# Spatial Attention Weakly Modulates Visual Responses in the Lateral Geniculate Nucleus

**DOI:** 10.1101/2025.05.16.652890

**Authors:** Henry J. Alitto, Jeffrey S. Johnson, W. Martin Usrey

## Abstract

Visual responses in the cerebral cortex are strongly influenced by shifts in spatial attention. This modulation of visual processing includes changes in firing rate, decreased response variability, and decreased interneuronal correlations; all of which are thought to underlie enhanced visual perception near the center of attention at the cost of visual perception at other locations. Visual information from the retina is relayed to primary visual cortex via neurons in the lateral geniculate nucleus (LGN) of the dorsal thalamus. Although early studies describe an enhancement of LGN activity with spatial attention, more recent work has cast doubt on this view. Given its strategic position as the gateway to the cortex, an understanding of the effects of attention on visual processing in the LGN is important. We therefore performed experiments to reexamine the influence of covert spatial attention on the spiking activity of single units in the macaque LGN and applied a broad set of analyses and functional metrics to assess possible effects. Our results reveal a statistically significant effect of spatial attention in the LGN: firing rates were slightly higher and more reliable when monkeys directed attention towards the receptive fields of recorded neurons compared to when attention was directed to different retinotopic locations. However, effects were much smaller than previously reported (∼1% vs ∼4%) and further analyses suggest that effects are weak and inconsistent. Thus, while spatial attention does exert an influence in the LGN, its effects are weak and may have limited impact on downstream processing.

**Significance Statement:** The lateral geniculate nucleus (LGN) is a critical relay in the visual system, shaping the flow of sensory information from the eye to the brain. Although higher-order brain regions show strong modulation by attention, it remains unclear whether the LGN is similarly affected. By directly recording LGN activity in monkeys performing a covert attention task, we found that spatial attention produces only weak and inconsistent modulation of visual responses. These findings suggest that the LGN largely operates independently of spatial attention, highlighting a potential boundary between early sensory processing and cognitive control.

## Introduction

Spatial attention refers to the allocation of perceptual resources to enhance the processing of visual information at a specific spatial location, improving stimulus detectability and behavioral responsivity (Posner, 1980). While a full understanding of spatial attention is still emerging, the neural correlates of volitional spatial attention are generally believed to involve top-down processes (Maunsell, 2015; Boshra and Kastner, 2022; Martinez-Trujillo, 2022). These processes modulate fundamental aspects of visual responses in the cerebral cortex, including changes in firing rate, decreased response variability, and decreased neuronal correlations (Luck et al., 1997; McAdams and Reid, 2005; Cohen and Maunsell, 2009; Mitchell et al., 2009). Consistent with a top-down framework, the influence of spatial attention on visual response properties is greatest towards the top of the visual hierarchy (e.g., IT cortex) and weakest in primary visual cortex (V1), the initial stage of visual processing in the cortex (Luck et al., 1997; Kastner and Ungerleider, 2000; McAdams and Reid, 2005; Lauritzen et al., 2010; Ni and Maunsell, 2019; Meyyappan et al., 2025).

Along with the cortex, subcortical areas also play a role in spatial attention (Saalmann and Kastner, 2011; Zénon and Krauzlis, 2012; Zhou et al., 2016; Sherman and Usrey, 2021). Two of these areas, the superior colliculus in the midbrain and the pulvinar in the thalamus, receive strong, driving input from the cortex (Lynch et al., 1985; Sherman and Guillery, 2002; Cerkevich et al., 2014; Baldwin et al., 2017) and, thus, a route exists for cortical mechanisms to influence their response properties and communication with target structures. In contrast, neurons in the lateral geniculate nucleus (LGN) lack driving cortical input, instead receiving their driving input exclusively from the retina which is then relayed to cortex (reviewed in Usrey & Alitto, 2015). Despite this, LGN activity has been reported to be influenced by spatial attention (O’Connor et al., 2002; McAlonan et al., 2008), potentially mediated by modulatory feedback pathways from the cortex and/or inhibitory circuits involving the thalamic reticular nucleus (TRN) (McAlonan et al., 2008). The impact of attention on LGN activity remains unclear, though, as more recent work has not found effects of attention on visual processing in the LGN (Jiang et al., 2015; Shah et al., 2022).

Given the differences in past results and the importance of understanding the dynamics of visual signals delivered to cortex, we reexamined the influence of spatial attention on the spiking activity of LGN neurons in the macaque monkey and applied a broad set of analyses and functional metrics to assess possible effects. Moreover, recognizing that effects may be small, we collected data from a large sample of LGN neurons to strengthen the statistical analyses. Our results reveal small but significant effects of attention on the firing rate and reliability of LGN responses to visual stimuli. However, a deeper analysis of the data reveals that these effects are weak and inconsistent. Specifically, we found that shifting spatial attention towards LGN receptive fields was the equivalent of increasing the stimulus contrast by 1.25% or 1.0% for parvocellular and magnocellular neurons, respectively. Likewise, an ROC analysis indicates that an ideal observer would only be able to distinguish attend-toward trials from attend-away trials with 52% accuracy. Additionally, we did not observe a change in burst activity with shifts in spatial attention. Given that changes in burst firing are considered a reliable indicator of changes in thalamic network state (Sherman, 2001; McCormick et al., 2015), this is further evidence of the weak influence of spatial attention on LGN responses. Finally, by applying a linear classifier to pseudo-populations constructed from our recorded sample of LGN neurons, we found that attentional state could indeed be decoded at the population level. However, consistent with our single-cell analysis, this signal was largely driven by a small subset of neurons with stronger attentional modulation, rather than emerging from a distributed population code. Collectively, these results indicate that the modulation of LGN responses by spatial attention is statistically significant but functionally ineffectual.

## MATERIALS AND METHODS

Two adult rhesus monkeys (Macaca mulatta; one male and one female) were used for electrophysiological recordings in this study. All experimental procedures conformed to the National Institutes of Health and the United States Department of Agriculture guidelines and were approved by the Institutional Animal Care and Use Committee at [**redacted for blind review**]. Under full surgical anesthesia, the animals received a cranial implant consisting of a head post for head stabilization and a recording cylinder that allowed access to the LGN.

### Electrophysiological recordings

Extracellular recordings from the LGN were made with platinum-glass electrodes (1–2 MΩ; Alpha Omega) using a microdrive (40 mm electrode travel; Thomas Motorized Electrode Manipulator, Thomas Recording) mounted on the recording chamber. Online continuous voltage signals were amplified (A-M Systems), filtered (0.1–5 kHz), and recorded using a Micro1401 data acquisition system (28 kHz) and Spike2 software (Cambridge Electronic Design, CED). To extract action potentials, extracellular signals were high-pass filtered (stopband edge frequency = 500 Hz), thresholded, and clustered within the Spike2 software.

### Visual Stimulation and Receptive Field Mapping

Visual stimuli were generated with a ViSaGe (Cambridge Research Systems) and presented on a gamma-corrected CRT monitor (Sony or Mitsubishi) positioned ∼80 cm in front of the animal. The display had a resolution of 1024×768, a refresh rate of 100-140Hz, and a mean luminance of ∼38 cd/m2. LGN receptive fields were manually mapped using small (<0.5°) user-controlled, computer-generated visual stimuli (custom software with Spike2 interface). The visual stimuli used for assessing spatial attention are described below.

### Behavioral training and performance

Data was collected while the animals performed a spatial attention task (Figure 1A-B) while eye position and pupil size were monitored using an Eye-Trac6 infrared eye tracker (Applied Science Laboratories) with a custom Spike2 interface. For this task, the animals initiated a trial by fixating a central fixation dot similar to the fixation dot used for passive fixation (hand mapping trials). The fixation dot was ∼0.2° diameter, and the fixation window was 0.8°-1.2° diameter. After a short delay (0.5-1.0s), two sine wave gratings (temporal frequency = 5 Hz, spatial frequency = 1.0 cycles/°) appeared at equidistant locations from the fovea, typically at an eccentricity of 4-8°. During recording sessions, one stimulus was positioned inside the recorded LGN receptive field (dashed black circle; note: the dashed circle was not present in the actual stimulus display presented to the animals). Each sine wave grating was surrounded by a colored circle (one green, one red), and the color of the circles at each location was constant throughout the task. The color of the fixation dot (either red or green) indicated to the animal which grating was 90% likely to change contrast (valid trials) after a delay with an exponential hazard function of 0.6 – 1.8s. On 10% of the trials (invalid trials), the uncued grating increased in contrast (same hazard function). Because one stimulus was always inside the RF and the other outside, trials were further categorized based on the spatial relationship between the cued location and the RF. “Attend toward” trials (A) occurred when the cued stimulus—i.e., the one most likely to change contrast—was positioned inside the RF. “Attend away” trials (B) occurred when the cued stimulus was positioned outside the RF. This classification was used in all subsequent analyses to assess how spatial attention modulated LGN responses. The base contrast of the grating was set to be near the C50 (contrast to evoke a half-maximum response) of the recorded LGN neuron, as estimated through hand mapping. After the contrast change, the animal was given 1s to respond by making a saccade to the location of the change, and the animal was rewarded with a small juice reward. Psychophysical performance (accuracy and reaction time) for valid and invalid trials were compared to ensure that the animals were applying spatial attention as anticipated. Based on these comparisons, the contrast change was titrated so that the animal performed 20-30% better on cued trials relative to uncued trials.

**Figure 1.**
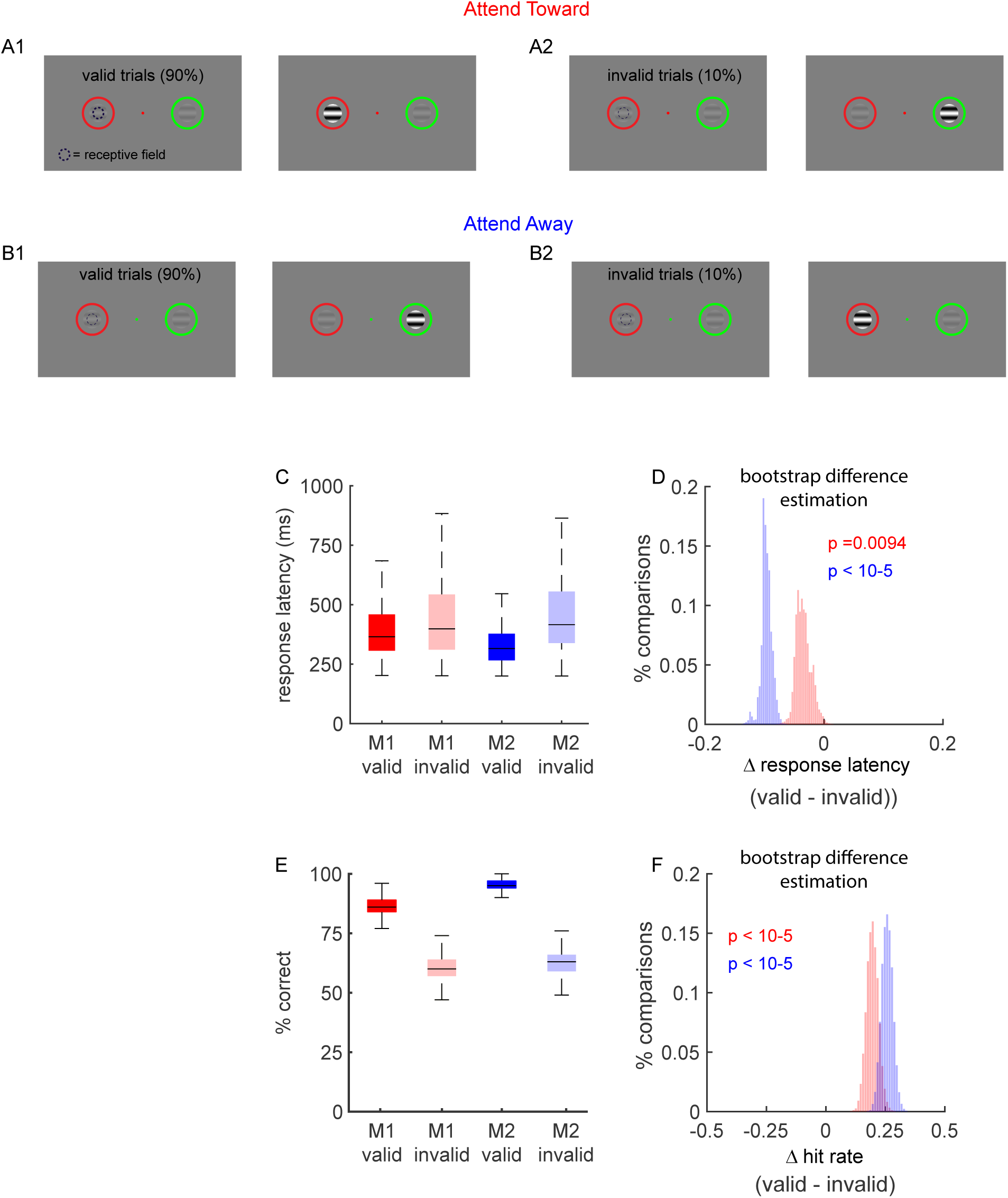
Spatial attention task and psychophysical measure of spatial attention. Two macaque monkeys (Monkey 1: M1 and Monkey 2: M2) were trained to perform a covert spatial attention task. ***A-B***. On each trial, animals maintained fixation on a central green dot while two peripheral sine wave gratings appeared, each surrounded by a colored circle. One grating was positioned over the receptive field of a recorded LGN neuron (dashed black circle) and the other at an equidistant location. The color of the fixation point, in conjunction with the colors of the peripheral circles, indicated the probability that each grating would undergo a contrast increase after a brief delay. In 90% of trials (**A1, B1**), termed “valid trials”, the contrast change occurred at the location where the circle color matched the fixation cue. In 10% of trials (**A2, B2**), termed invalid trials, the contrast change occurred at the non-matching location. Since one stimulus was positioned inside the receptive field and one was positioned outside the receptive field, trials were further categorized based on the spatial relationship between the cued location and the RF. “Attend toward” trials **(A)** occurred when the cued stimulus was positioned inside the receptive field, while “attend away” trials **(B**) occurred when the cued stimulus was positioned away from the receptive field. The error bars in **C and B** indicate the 25^th^ and 75^th^ percentiles. Animals were rewarded for making a saccade to the grating that increased in contrast. Both animals respond faster (***C, D***) and more accurately (***E, F***) on valid trials compared to invalid trials. Statistical significance was assessed using bootstrap difference estimation (BDE). Panels D and F show the distribution of median differences (valid – invalid) computed over 10,000 bootstrap resamples for M1 (red) and M2 (blue). These distributions provide graphical estimates of effect size, variability, and statistical confidence (see Methods and Materials for details).

#### Analysis

##### Classification of neurons

Based on the hand-mapping procedures described above, we classified LGN neurons on two axes: cell-type and response polarity (Alitto et al., 2011). Cell types were defined as magnocellular or parvocellular based on the C50 (C50 greater than 35% were classified as parvocellular, C50 less than 35% were classified as magnocellular). Response polarity (on-center cells vs off-center cells) was based on responses to increases and decreases in luminance.

##### Firing rate and spatial attention

LGN neurons have linear responses to the presentation of drifting sine wave gratings that are modulated by the phase of visual stimulus. To extract this linear response, we calculated the average cycle histogram (cycle duration = 200 ms, bin size = 7.7 ms) for each trial and extracted the linear response as the F_1_ from a Fourier transform with a Hanning window (custom Matlab script). Based on the F1 and F0 (mean firing rate), the attention index was defined as:

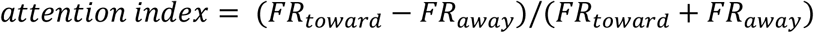

Here, the FR is the firing rate (either the F_0_ or F_1_) during attend toward and attend away trials.

##### Equivalent Contrast Change

Shifts in spatial attention are often analogized as being similar to changes in stimulus contrast (Carrasco et al., 2004; Reynolds and Chelazzi, 2004). Given that shifts in spatial attention cause changes in the firing rates of visual neurons, we wanted to determine the changes in contrast that would have produced equivalent changes in firing rate. In the current study, the baseline stimulus contrast was individually specified for each LGN neuron, based on computer-aided hand mapping of the receptive field, to evoke 50% of the cell’s maximum response (C50). Since complete contrast response functions were not available for the current data set, we created standardized parvocellular and magnocellular contrast response functions from a separate sample of LGN neurons (Figure 3a; magnocellular: n = 33, parvocellular: n = 87; 5-8 stimulus repeats per contrast) and applied the attentional changes in firing onto this value to determine an effective/equivalent contrast change. For each recorded neuron, we estimated the attention-driven change in firing rate as:

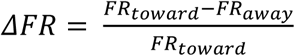

**Figure 2.**
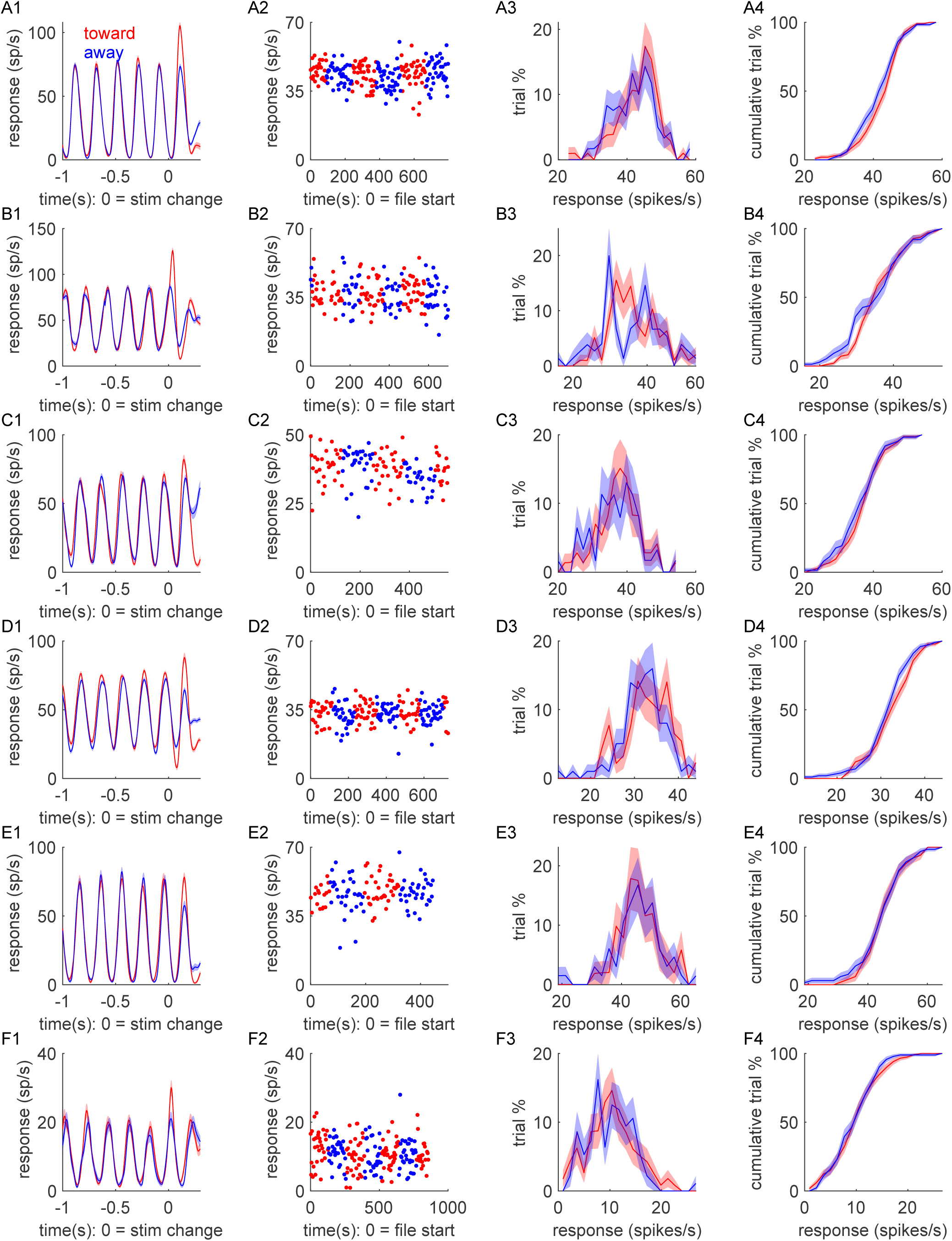
Single-unit examples of lateral geniculate nucleus (LGN) responses recorded while the animal performed the spatial attention task. Each row (***A - F***) shows data from one neuron. ***Column 1***. Peri-stimulus time histograms (PSTHs) showing firing rates for valid *attend-toward* (red) and *attend-away* (blue) trials, aligned to the time of contrast change (0 s). ***Column 2***. Mean firing rate across all trials as a function of time relative to the start of recording. ***Column 3***. Distribution of firing rates (F_1_) across all trials. ***Column 4***. Cumulative distribution of firing rates across all trials. In all panels, the shaded areas indicate standard error of the mean.

**Figure 3.**
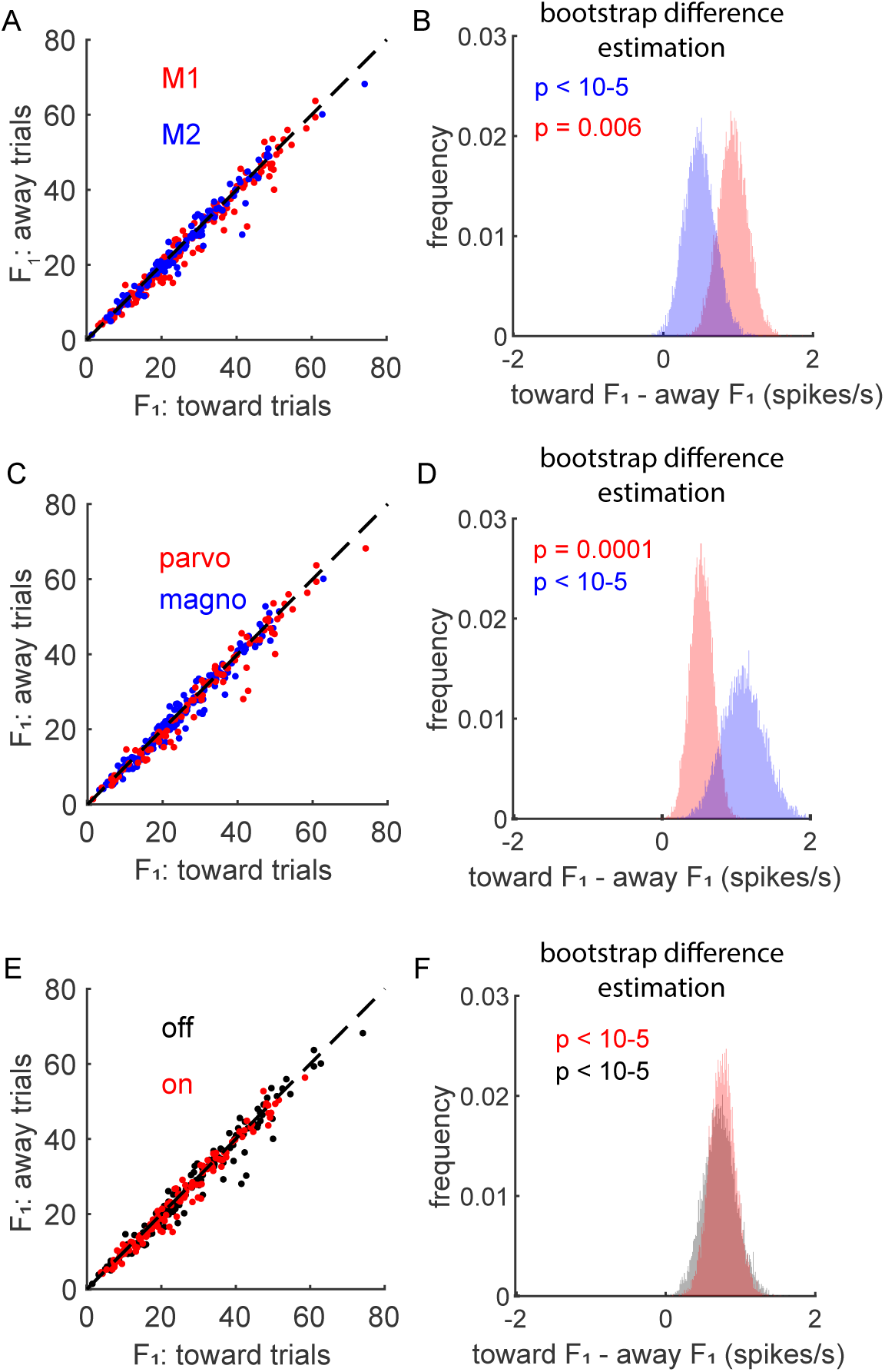
Firing rate during spatial attention. ***A***. Distribution of firing rates (f1) for attend toward vs attend away trials across all cells (M1 = red, M2 = blue; n = 283). ***B***. Significance of these effects was determined via a bootstrap difference estimation (See Materials and Methods). The red and blue data in panel ***B*** show the distribution of median difference values (toward – away) over 10,000 resamples with replacement for M1 and M2, respectively. These distributions provide a direct graphical illustration of statistical significance and confidence intervals (p values in text). ***C, D***. Same as ***A, B*** for parvocellular (red) and magnocellular (blue) data. ***E, F***. Same as ***A, B*** for on-center (red) and off-center (black).

Using the standardized magnocellular and parvocellular contrast response functions, we then estimated the contrast that would have evoked an equivalent response difference relative to the C50. For example, if the attend-toward firing rate was 1.2x greater than the attend-away firing rate, we would identify the contrast that would have evoked a response that was 1.2x the firing rate evoked at C50. Based on this calculation we then defined attention equivalent contrast change as:

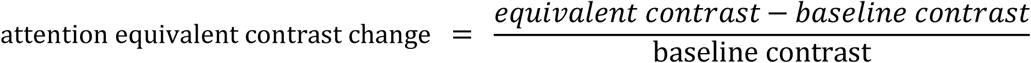

##### ROC analysis of attentional modulation

To further quantify the influence of spatial attention on visual responses in the LGN, we performed an ROC analysis to determine how well an ideal observer would be able to determine the focus of the animal’s attention based solely on the trial-by-trial distribution of F_1_s across attend-toward and attend-away trials. Using the matlab function perfcurve (bootstrap n = 1000), we determined if (a) attend toward trials were distinguishable from spontaneous activity, (b) attend away trials were toward trials were distinguishable from spontaneous activity, and (c) attend toward trials were distinguishable from attend away trials. For this analysis, spontaneous activity was based on the intertrial interval (ITI).

##### Influence of attention on thalamic Bursts

Spiking activity in the thalamus is often segregated into tonic and burst firing (Sherman, 2001). In tonic firing, thalamic neurons respond to excitatory stimuli in a linear, constant fashion where the mean firing rate is proportional to the strength of the stimulus. When thalamic neurons are sufficiently hyperpolarized to deinactivate T-type Ca^++^ channels, they begin to exhibit bursting activity (McCormick and Huguenard, 1992). As has been done previously (Reinagel et al., 1999; Alitto et al., 2019; Sanchez et al., 2023), we tracked burst and tonic firing based on the following criteria: (1) an ISI of >100 ms that preceded the first spike in a sequence, and (2) one or more subsequent spikes that followed with ISIs of <4 ms. Past studies applying these criteria to intracellular recordings show that events defined as bursts co-occur with T-channel plateau potentials (Lu et al., 1992). Based on these criteria, we estimated the modulation of thalamic burst firing in the LGN by spatial attention.

##### Modulation of response reliability by spatial attention

To quantify the influence of spatial attention on response reliability in the LGN, we calculated the Fano factor for both attend-toward and attend-away trials (custom Matlab function). Fano factor is a common measure of response reliability and is defined as:

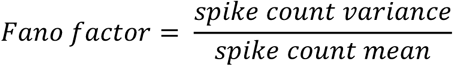

##### Support vector machine

To quantify the influence of spatial attention at the population level, we trained linear support vector machine (svm) classifiers using MATLAB’s fitcsvm function on pseudo-populations of sequentially recorded LGN neurons (Hastie et al., 2009). Pseudo-populations were constructed using two strategies: with and without cell identity, both incorporating bootstrap sampling over 10,000 iterations.

In the with cell identity condition, we constructed pseudo-population trials by randomly sampling (with replacement) 20 attend-toward and 20 attend-away trials from each neuron. This resulted in feature vectors with 198 features (one per neuron) and 40 trials per bootstrap iteration. In the without cell identity condition, we generated feature vectors of the same size (198 features × 40 trials), but the firing rates were drawn randomly from the entire pool of trials, independent of neuron identity—effectively destroying the mapping between neurons and features.

In both approaches, the SVM was trained on the pseudo-population and tested on a separate, non-overlapping validation set constructed using the same sampling procedure. Classifier performance was quantified using MATLAB’s predict and perfcurve functions, with performance defined as the area under the curve (auc, receiver operator characteristic) on the validation data. To establish a chance-level baseline, we implemented a shuffle control in which trial labels (attend-toward vs. attend-away) were randomized prior to validation. Shuffled-label decoding was performed using the same bootstrap framework. Statistical comparison between real and shuffled models was based on the proportion of bootstrap iterations in which the real-model AUC exceeded the shuffled AUC.

To assess the distribution of attentional modulation across the LGN population, we implemented two additional SVM-based analyses: greedy ablation and reverse greedy ablation. In the greedy ablation analysis, we began with the full pseudo-population and iteratively removed the most informative neuron—defined as the neuron with the largest absolute SVM weight (beta coefficient). In the reverse greedy ablation analysis, we removed the least informative neuron at each step. In both cases, the classifier was retrained and evaluated after each ablation step using the same procedure described above.

##### Statistical tests

When testing for statistical differences between two populations, we performed bootstrap analyses to estimate median values, median difference values, 95% confidence intervals, and statistical significance. For this process, data were resampled with replacement 10,000 times while maintaining the original sample size in each resampling iteration. Within the text, statistics will be presented as:

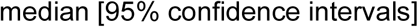

> for example:

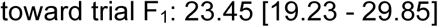

Where applicable, a bootstrap median difference estimation (BDE) is also provided. Throughout the Results, the distributions from bootstrap analyses are illustrated within the figures (e.g., Figure 1D-F). These histograms show the distribution of median values (or median difference values) generated from the 10,000 resamples with replacement. These distributions provide graphical estimates of effect size, variability, and statistical confidence. For within-neuron statistical tests, trials were shuffled across attention conditions (e.g., attend toward vs. attend away) and resampled with replacement 10,000 times per neuron. Statistical significance for each neuron was determined as the proportion of resampled values exceeding the observed effect size:

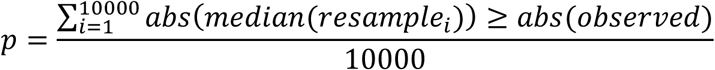

For sample-level statistical tests involving index values (e.g., attention index, burst index), significance was defined as the proportion of resampled values having the different sign as the observed value. Specifically, for an observed attention index >0, significance was calculated as:

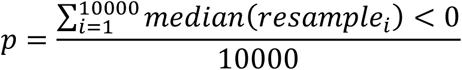

For an observed attention index <0, significance was calculated as:

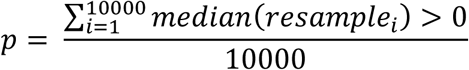

## Results

To study the influence of attention on visual processing in the lateral geniculate nucleus (LGN), we trained two macaque monkeys (M1 and M2) to perform a spatial attention task (Figure 1A-B, see Materials and Methods). Briefly, the animals were trained to fixate on a colored dot while two drifting sinusoidal gratings, each surrounded by a thin colored circle, were shown in the periphery (mean eccentricity = 7.7°±3.3°, std). The contrast of the drifting gratings was held constant for a randomized interval (0.6s – 2.0s), at which point one of the gratings increased in contrast and the animal was rewarded for making a saccade to that location. The probability of each grating changing contrast was linked to the color of the fixation point. If the color of the fixation point matched the color of the surrounding circle, then that grating changed contrast on 90% of the trials (valid trials). On the remaining 10% of trials, the nonmatching grating changed contrast (invalid trials). Importantly, the color of the circles did not change from trial to trial; instead, the color of the fixation point changed. As expected, both monkeys were faster and more accurate on valid trials relative to invalid trials, indicating the animals were using covert spatial attention to perform the task (Figure 1 C-F; response latency: validM1 = 354ms [347 - 365], invalid_M1_ = 390ms [361 - 405], BDE_M1_ = -34ms [-54 - -5], p_M1_ = 0.0094; valid_M2_ = 301ms [298 - 303], invalid_M2_ = 396ms [378 - 417], BDE_M2_ = -96ms [-117 - -77], p_M2_ < 10^-5^); hit rate: valid_M1_ = 0.949 [0.932 - 0.964], invalid_M1_ = 0.749 [0.701 - 0.796], BDE_M1_ = 0.20 [0.152 - 0.249], p_M1_ < 10^-5^; valid_M2_ = 0.982 [0.979 - 0.985], invalid_M2_ = 0.719 [0.672 - 0.767], BDE_M2_ = 0.263 [0.217 - 0.311], p < 10^-5^).

To quantify the influence of spatial attention on the spiking activity of LGN neurons, we recorded the responses of 283 isolated single units (Figure 2, 103 magnocellular neurons, 180 parvocellular neurons; 151 off-cells, 132 on-cells) across both monkeys while they performed the spatial attention task. For each neuron, one of the drifting gratings was centered over the neuron’s receptive field and the other grating was shown at an equidistant location from the fixation point. Gratings were approximately 2-3 times the estimated size of the recorded neuron’s classical receptive field (typically 2° in diameter). Across cells and illustrated with the examples shown in Figure 2, LGN responses to the drifting sine wave grating largely matched the drift frequency of the grating (5 Hz). Thus, as is standard, we primarily focused on this component (first harmonic, F_1_) of the firing rate as quantified through Fourier analysis (see Materials and Methods).

### Influence of spatial attention on neuronal firing rate

Across our sample of LGN neurons, there was a small, but significant, increase in the F1 of the firing rate when the monkey directed attention toward the LGN cell’s receptive field (toward trials) relative to when it directed attention away from the receptive field (away trials) (Figure 3). This effect was seen in M1 (Figure 3A-B (red data), toward = 26.1 [23.9 - 28.4] spikes/s, away = 25.2 [22.9 - 27.5] spikes/s, n = 161, BDE = 0.93 [0.56 - 1.32] spike/s, p < 10^-5^) and M2 (Figure 3A-B (blue data), toward = 25.8 [23.1 - 28.4] spikes/s, away = 24.9 [22.255 - 27.476], n = 122, BDE = 0.48 [0.10 - 0.92] spikes/s, p = 0.0059). This effect was also seen when data were divided based on cell type (Figure 3C,D (parvocellular, red data), toward = 24.5 [22.8 – 26.2] spikes/s, away = 23.9 [22.2 – 25.7] spikes/s, BDE = 0.54 [0.24 – 0.84] spike/s, p = 0.0001; (magnocellular, blue data), toward = 29.1 [26.1 – 32.1] spikes/s, away = 28.0 [25.0 – 31.0], BDE = 1.1 [0.57 – 1.67] spikes/s, p < 10^-5^) and luminance polarity (Figure E-F, (off-center, black data) toward = 26.9 [24.7 – 29.2] spikes/s, away = 26.2 [24.0 – 28.5], BDE = 0.72 [0.31 – 1.17] spikes/s, p < 10^-5^; (on-center, red data), toward = 25.2 [ 23.0 – 27.4] spikes/s, away = 24.5 [22.4 – 26.7] spikes/s, BDE = 0.76 [0.43 - 1.1] spikes/s, p < 10^-5^). The differences between parvocellular vs magnocellular (BDE = -0.16 [-0.73 – 0.33] spike/s, p = 0.31) and on-center vs off-center (BDE = 0.12 [-0.36 – 0.60] spikes/s, p = 0.65) were not statistically significant. Similar differences were seen for the mean response (F0; data not shown), for M1 (toward = 27.4 [24.9 - 30.0] spikes/sec, away = 26.5 [24.0 - 29.0] spikes/sec, BDE = 0.92 [0.54 - 1.32] spikes/s, p < 10^-5^) and M2 (toward = 30.6 [27.7 - 33.4] spikes/s, away = 30.0 [27.2 - 32.9] spikes/s, BDE = 0.54 [0.06 - 1.08] spikes/s, p < 0.0114). Across monkeys and cell types, the influence of attention on firing rate was significant, but small, increasing firing rate by ∼ 1 spike/s.

Given that the firing rate varies from cell to cell, a bounded index is often useful for making comparisons between conditions across a sample of cells. We therefore used an attention index based on each cell’s F_1_ response, where

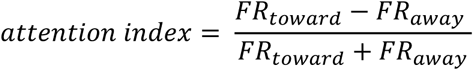

With the attention index, positive values indicate the firing rate increased with attention directed towards the neuron’s receptive field and negative values indicate the firing rate decreased with attention. For both monkeys, median attention index values were significantly greater than zero (Figure 4A, D-E; M1: 0.014 [0.011 - 0.022], p < 10^-5^; M2: 0.01 [0.001 - 0.012], p = 0.0009). When data was combined across monkeys an increased attention index was evident for parvocellular neurons (Figure 4F-G, 0.011 [0.003 - 0.015], p = 0.0004, n = 180), magnocellular neurons (Figure 4F-G, 0.012 [0.009 - 0.020], p < 10^-5^, n = 103), on-cells (Figure 4H-I, 0.012 [0.007 - 0.018], p < 10^-5^, n = 131), and off-cells (Figure 4H-I, 0.012 [0.004 - 0.015], p < 10^-5^, n = 152). The differences between parvocellular and magnocellular neurons (parvocellular – magnocellular: -0.002 [-0.012 - 0.004], p = 0.22) as well as between on-center and off-center neurons (on-center – off-center: 0.001 [-0.006 - 0.009], p = 0.41) were not statistically significant. On a cell-by-cell basis, 16.6% of cells (47/283) had a statistically significant attention index at a significance level of 0.05 (Figure 4B). This distribution of statistically significant cells was marginally biased (Figure 4C, p = 0.0479) toward significantly positive attention index values (11.1% of cells) compared to significantly negative attention values (5.3% of cells). For the remainder of the paper, no statistically significant differences were seen between on-center and off-center data. Thus, on-center vs off-center differences will not be discussed.

**Figure 4.**
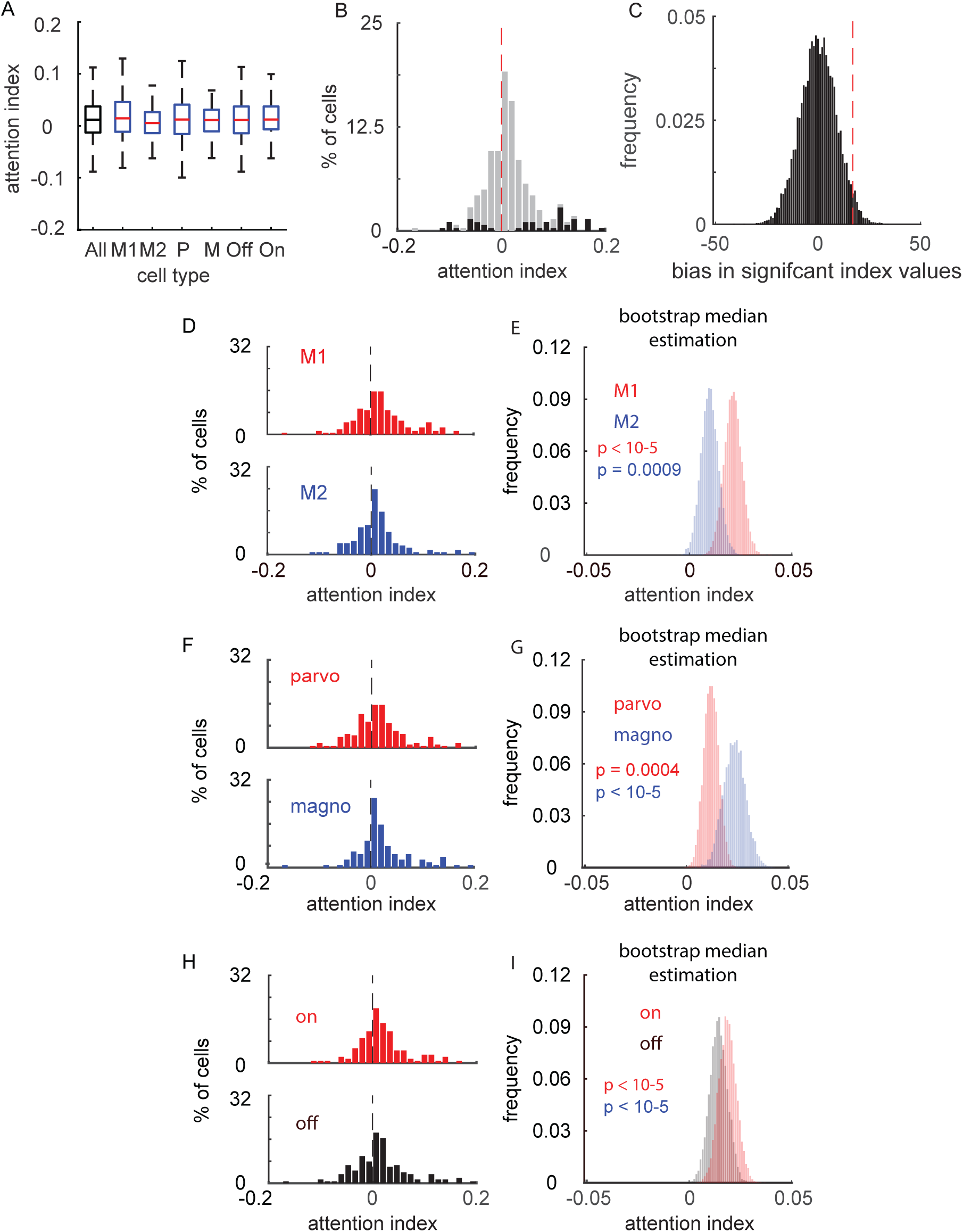
Attention index. ***A***. Distribution of attention index values across cell types (error bars = 25^th^ and 75^th^ percentiles). ***B***. Overall distribution of attention index values. Significant values (p< 0.05) are indicated in black. ***C***. Significance of the distribution of attention index values (positive > negative) was determined by a bootstrap null distribution analysis (red dashed line = observed difference). ***D***. Distribution of attention index values for M1 (red) and M2 (blue) (0 = black dashed line). ***E***. Significance of these distributions was determined via a bootstrap mean estimation. ***F, G***. Same as ***D, E*** for parvocellular (red) and magnocellular (blue) data. ***H, I***. Same as ***A, B*** for on-center (red) and off-center (black) data.

Given that the influence of attention on firing rates appeared to be relatively small, we wished to put this change into perspective and estimate its functional relevance. Since spatial attention is often viewed as effectively increasing the contrast of the visual stimuli at that attended location, we wanted to determine the change in stimulus contrast that would evoke the same change in firing rate as shifting spatial attention toward the LGN receptive field (Figure 5). We therefore calculated the attention equivalent contrast change (AECC, see Materials and Methods). In the current study, the baseline contrast was set to be at the neuron’s C50 (the contrast that evoked a half-maximum response), but since each neuron’s C50 was estimated online with hand-mapping procedures, we used data from a separate population of neurons to generate standardized contrast response functions for magnocellular (n = 33) and parvocellular (n = 87) neurons (Figure 5A). Using these standardized functions, we calculated the contrast, relative to the C50, that evoked the same change in firing rate as shifting spatial attention. For example, if the change in firing rate caused by shifting attention was equivalent to the change in firing rate caused by increasing the contrast from 30% to 40%, then the AECC would be 0.33. For the parvocellular neurons in this study (Figure 5B-D), the 0.52 [0.097 - 0.785] spikes/s increase in firing when spatial attention was shifted toward the cell’s receptive field is equivalent to a relative increase in contrast of only 0.025 [0.017 - 0.032] that is a change equal to 2.5% of the C50. For a typical parvocellular neuron, this would be the equivalent of changing the contrast from 50.0% to 51.25%. For magnocellular neurons (Figure 5B-D), the 0.681 [0.294 - 1.112] spikes/s increase in firing rate with attention is equivalent to a relative increase in contrast of 0.058 [0.041 - 0.076]. For a typical C50 seen in magnocellular neurons, this would be the equivalent of changing the contrast from 18.0 to 19.0. Across cells, equivalent contrast change values were significantly greater for magnocellular neurons than for parvocellular neurons (magno - parvo: 0.033 [0.014 - 0.052] p = 0.0002); however, for both cell types the equivalent contrast change values were very small and likely would be below threshold for change detection.

**Figure 5.**
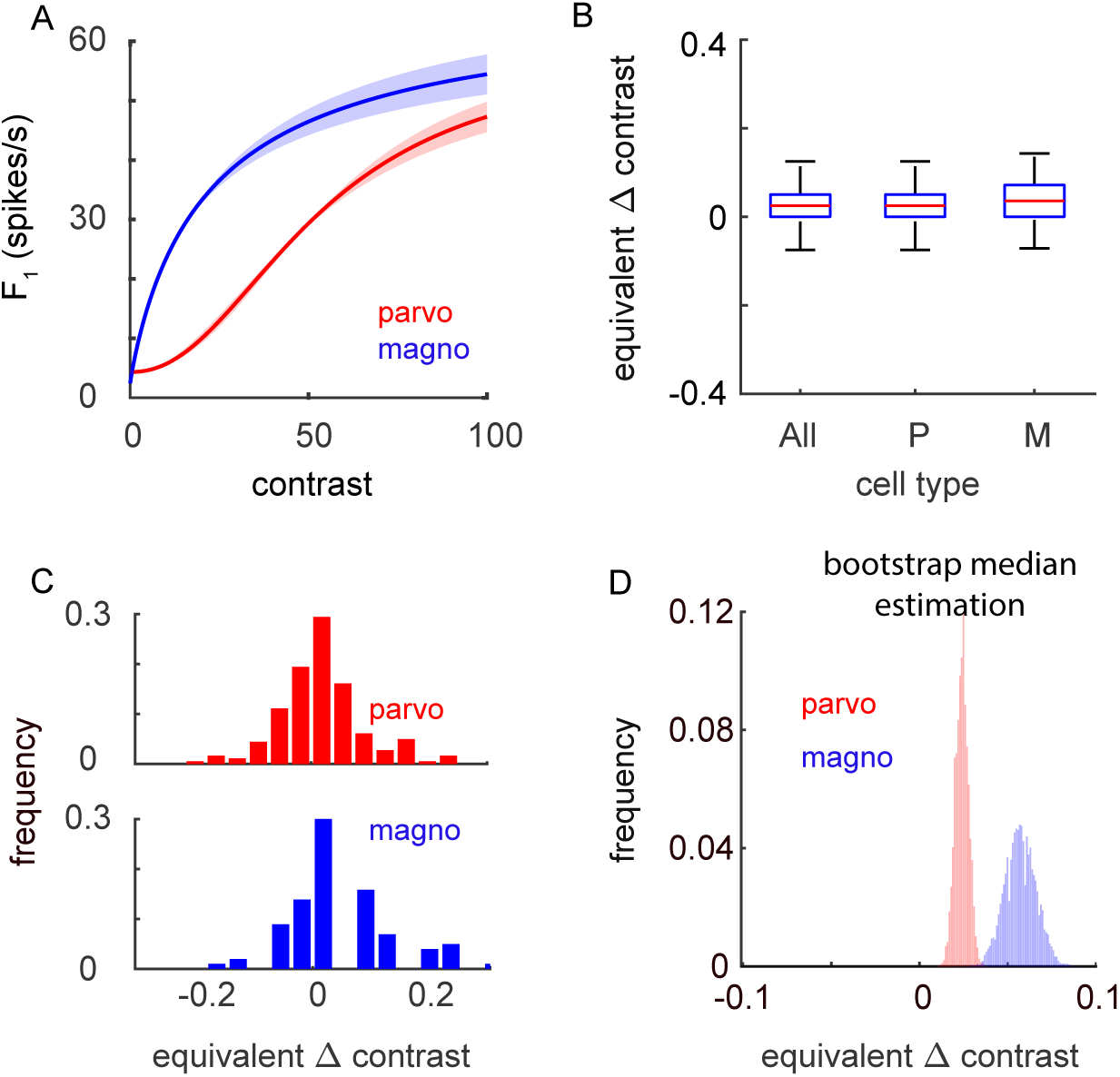
Equivalent contrast change. ***A***. Population contrast response functions for parvocellular (red) and magnocellular (blue) data (shaded area = standard error of the mean). ***B, C***. Distribution of equivalent contrast values across cell types (error bars = 25^th^ and 75^th^ percentiles). ***D***. Significance of these distributions was determined via a bootstrap mean estimation.

A separate metric that can be applied to assess the influence of spatial attention on visual responses utilizes receiver operating characteristic (ROC) analysis (Figure 6). With ROC analysis, one can ask how well an ideal observer would be able to determine, on a cell-by-cell basis, where the monkey was focusing its attention based solely on the response distributions (see Materials and Methods). A small difference in mean response would be discernable if there was also little variability (e.g., all trials with attention towards the cell’s RF had the same small difference in firing rate compared to all away trials). At the opposite extreme, the same small difference in mean response would not be discernable if associated with much greater variability. In the first case with low variance, the ideal observer would successfully distinguish attend-towards and attend-away trials a high percent of the time. In the second case, the ideal observer would do much worse. Applying this approach, results indicate an ideal observer would have no difficulty discriminating either toward or away trial contrast changes from the baseline contrast (Figure 6A-B, D-I, toward = 0.912 [0.898 - 0.925], p < 10^-5^; away = 0.902 [0.888 - 0.916], p < 10^-5^). On a cell-by-cell basis, 97.2% of cells had a statistically significant toward trial vs baseline AUC and 96.5% of cells had a statistically significant away trial vs baseline AUC (p<0.05). However, and more importantly, the same methodology revealed that toward and away trials could only be discriminated with an accuracy of 0.522 (Figure 6C, 0.522 [0.513 - 0.532]). On a cell-by-cell basis, only 13.8% of cells had statistically significant toward vs away AUC (p<0.05). However, it is worth nothing that significant AUCs were more common (p = 0.01) for cells with greater activity while attending toward the LGN RF (n = 28, 9.9% of cells) compared to cells with greater activity while attending away from the LGN RF (n = 11, 3.9% of cells). Overall, these results indicate the influence of spatial attention in the LGN is weak and not a reliable indicator of where the animal was attending on a trial-to-trial basis.

**Figure 6.**
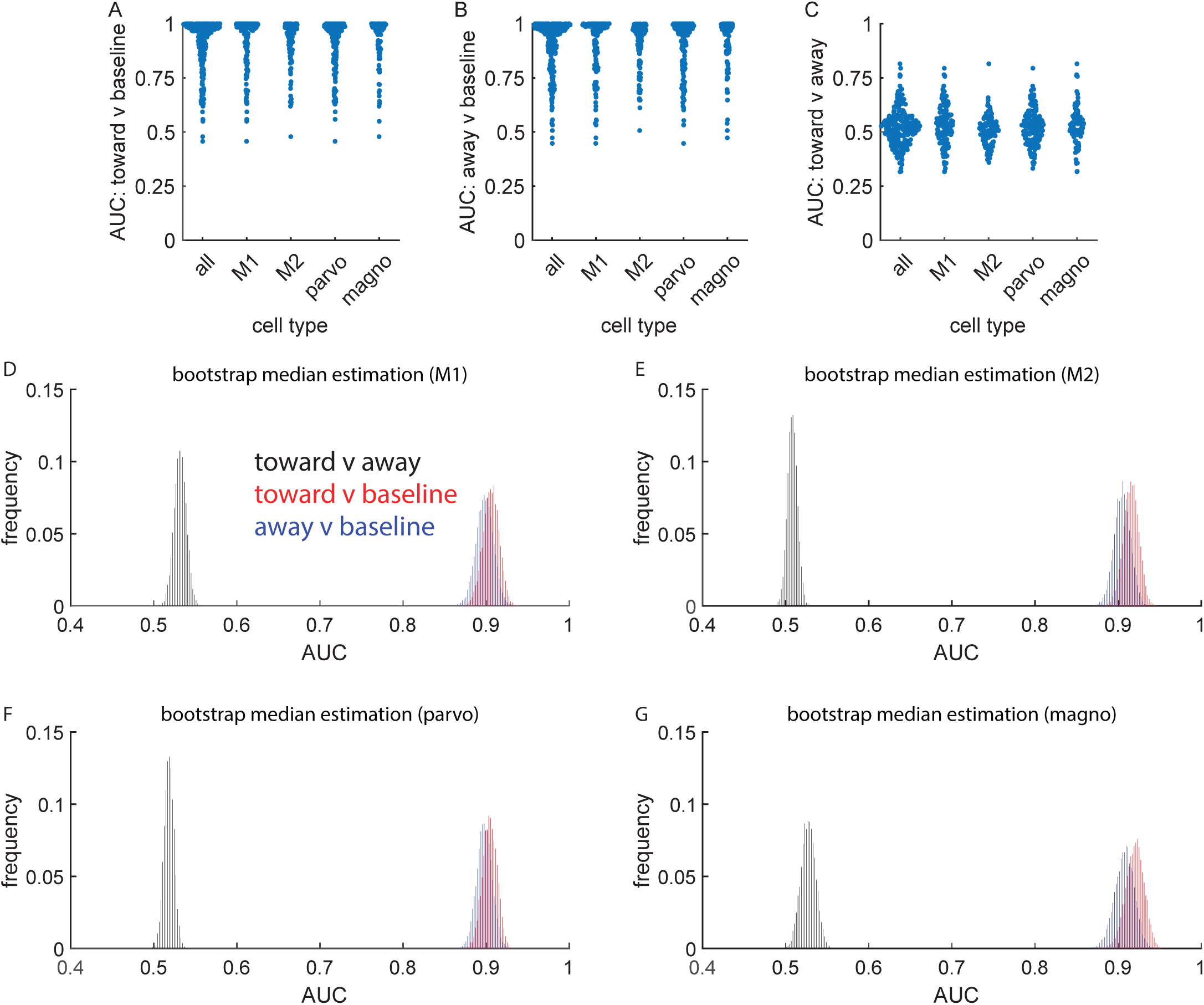
ROC analysis of spatial attention. ***A***. Area under the curve (AUC) for attend toward trials vs baseline, across cell types. ***B***. Area under the curve for attend away trials vs baseline, across cell types. ***C***. Area under the curve for attend toward trials vs attend away trials, across cell types. ***D***. Distribution of AUC values for M1 (red = toward vs baseline, blue = away vs baseline, black = toward vs away). ***E***. Same as ***D*** for M2. F. Same as ***D*** for parvocellular data. ***G***. Same as ***D*** for magnocellular data.

### Spatial attention and thalamic bursts

Across thalamic nuclei, relay cells produce spikes that can be categorized as burst spikes or tonic spikes (Sherman, 2001). Although burst spikes are less common in alert animals compared to anesthetized animals (Glenn and Steriade, 1982; Ramcharan et al., 2000; Alitto et al., 2011), spatial attention could serve to modulate the relative occurrence of burst and tonic spikes. Consistent with previous studies, results from this study show that burst spikes were infrequent in the alert animal (Figure 7A). This was the case for trials when animals were directed to attend towards the recorded cells’ receptive fields (proportion burst spikes: all cells = 0.017 [0.015 - 0.018], M1 = 0.018 [0.016 - 0.021]; M2 = 0.014 [0.011 - 0.017], parvocellular cells = 0.018 [0.015 - 0.020], magnocellular cells = 0.016 [0.013 - 0.019]) and away from the recorded cells’ receptive fields (proportion burst spikes: all cells = 0.016 [0.015 - 0.018]; M1 = 0.018 [0.016 - 0.021]; M2 = 0.015 [0.012 - 0.017], parvocellular cells = 0.017 [0.015 - 0.020]; magnocellular cells = 0.016 [0.013 - 0.019]). Further, shifts in spatial attention did not significantly modulate the probability of an LGN neuron firing in burst mode as assessed using a burst index:

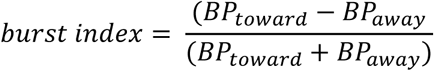

**Figure 7.**
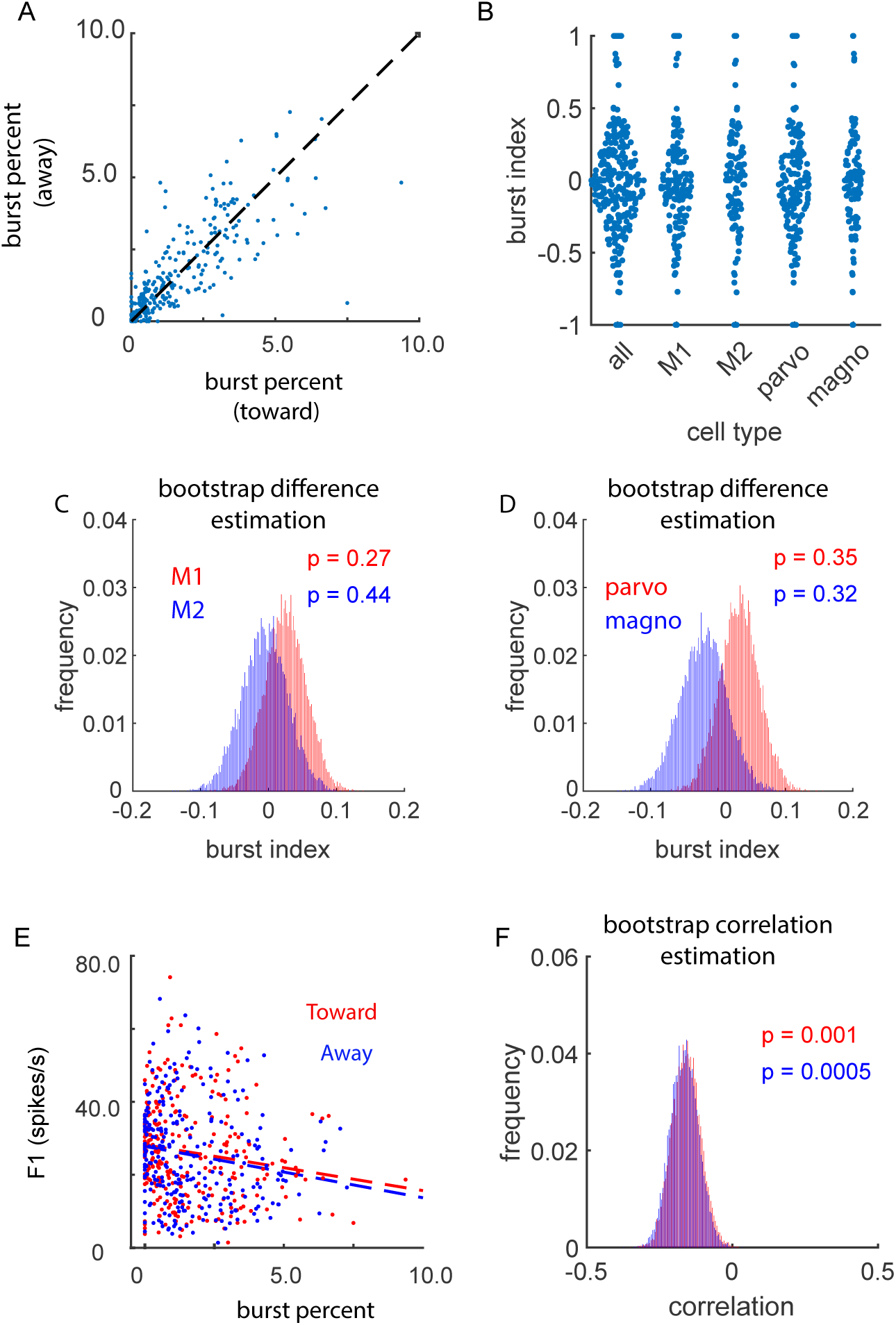
Thalamic bursts during spatial attention. ***A***. Distribution of thalamic burst percentage for attend toward trials vs attend away trials. ***B***. Sample distributions for burst index values across cell types. ***C, D***. Significance of these effects illustrated via a bootstrap difference estimation. ***E***. Correlation between burst percentage and firing rate across all cell types (red = attend toward trials, blue = attend away trials, dashed line = linear fit). ***F.*** Significance of these correlations illustrated via a bootstrap mean estimation.

(Figure 7B-D; burst index: M1 = 0.027 [-0.034 - 0.0088], p = 0.27; M2 = -0.004 [-0.074 - 0.061], p = 0.44; parvocellular cells = -0.017 [-0.072 - 0.029], p = 0.35; magnocellular cells = -0.009 [-0.088 - 0.009], p = 0.32). On a cell-by-cell basis, 11.7% of cells (33/283) had significant burst index values, evenly split between more bursts during attend-toward trials (5.6%) and more bursts during attend-away trials (6.0%) at a significance level of 0.05. There was, however, a significant negative correlation between the firing rate of recorded cells and burst percent for both toward trials (Figure 7E,F; r = -0.15 [-0.247 - -0.055], p = 0.001) and away trials (r = -0.17 [- 0.259 - -0.070], p = 0.0005). This inverse relationship might be explained by the time-voltage relationship that governs the production of bursts which, in turn, may reflect variable levels of alertness across/within recording sessions. Importantly, there was no correlation between burst rate and the magnitude of spatial attention effects on the firing rate of LGN cells (r = 0.03 [- 0.150 - 0.091], p = 0.31).

### Influence of spatial attention on response reliability

Past studies have shown that spatial attention increases response reliability in visual cortex (Mitchell et al., 2007); however, this effect has not been previously demonstrated in the LGN. To examine the influence of spatial attention on LGN response reliability, we calculated the Fano factor (Fano factor = spike count variance / spike count mean) when animals were directed to attend towards and away from the recorded neurons’ receptive fields (Figure 8). Consistent with results from studies not examining attention (Kara et al., 2000; Alitto et al., 2011), Fano factors across our sample of LGN neurons were, on average, below 1.0, indicating the responses of LGN neurons are less variable than expected from a Poisson process. This was the case for trials when animals were cued to attend towards the recorded cells’ receptive fields (Figure 8A, Fano factor: M1 = 0.928 [0.908 - 0.958]; M2 = 0.924 [0.897 - 0.951], parvocellular cells = 0.932 [0.900 - 0.955]; magnocellular cells = 0.922 [0.908 - 0.968]; and away from the recorded cells’ receptive fields (Fano factor: M1 = 0.92 [0.91 - 0.94], M2 = 0.95 [0.90 - 0.98], parvocellular cells = 0.932 [0.900 - 0.955]; magnocellular cells = 0.922 [0.908 - 0.968]). Although Fano factors were extremely similar for attend towards and attend away trials, there was a significant decrease in the Fano factor (as assessed using a Fano factor index, see Materials and Methods) when attention was directed towards the receptive fields of recorded neurons (Figure 8B-D); FF Index: M1 = -0.005 [-0.010 - 0.000], p = 0.0306; M2 = -0.008 [-0.013 - -0.003], p = 0.0003; parvocellular cells = -0.004 [-0.008 - 0.000], p= 0.036; magnocellular cells = -0.010 [-0.016 - -0.004], p = 0.0005). On a cell-by-cell basis 21.9% of cells had statistically significant Fano factor index values (62/283), at a significant level of 0.05, with a significant bias (p = 0.0002) toward more cells with significantly negative FF index values (18%, more reliable during attend-toward trials) compared cells with significantly positive FF index values (4.6%, more reliable during attend away trials). Thus, similar to the effects of attention on firing rate, there was a small, but significant, influence of attention on the response reliability of LGN neurons.

**Figure 8.**
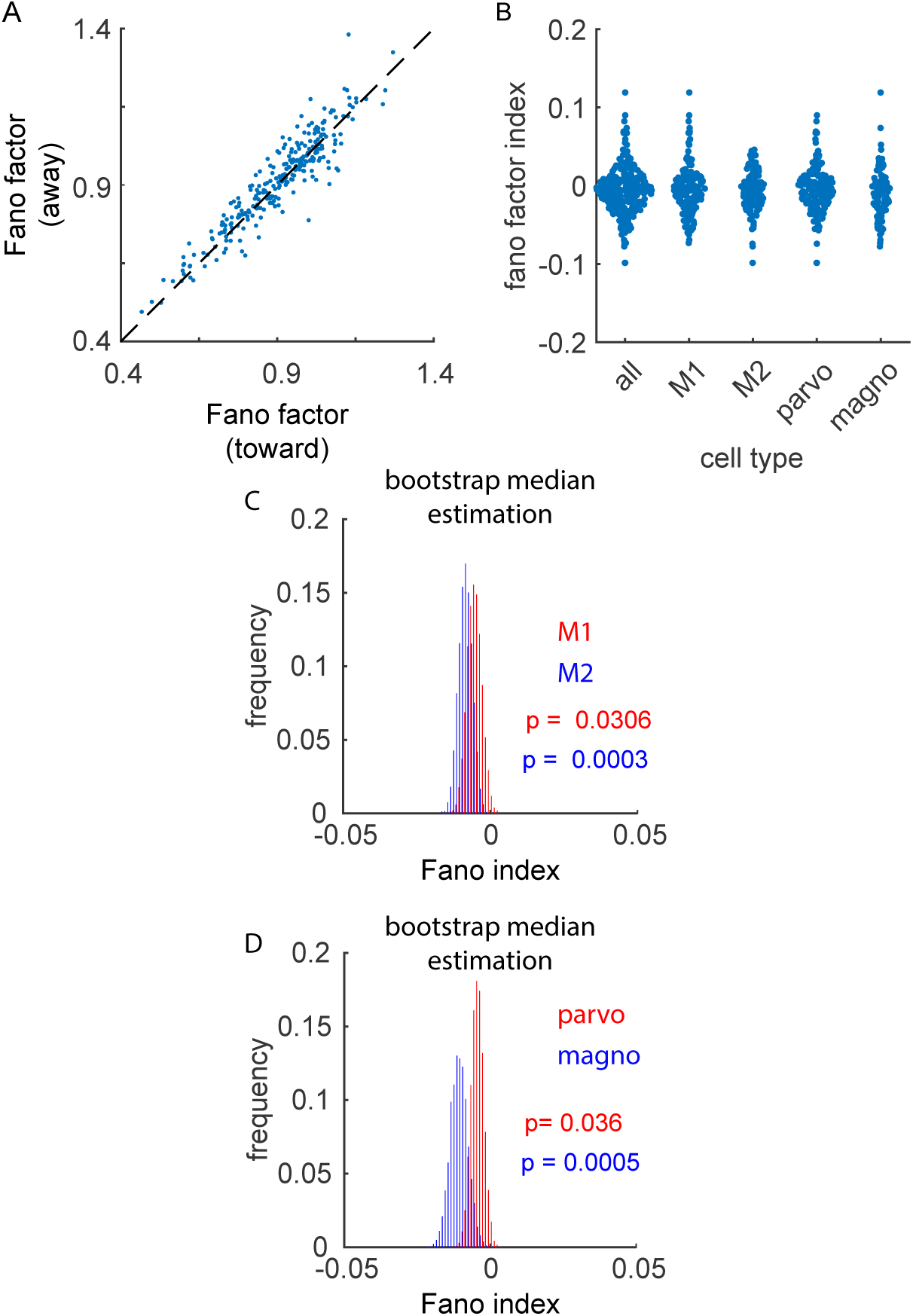
Response reliability during spatial attention. ***A***. Distribution of Fano factor values for attend toward trials vs attend away trials. ***B***. Sample distributions for Fano factor index values across cell types. ***C, D***. Significance of these effects as illustrated via a bootstrap difference estimation.

### Linear classification of LGN population activity

Although the influence of spatial attention on visual responses in the LGN is statistically significant, we have provided several lines of evidence that suggest the influence is unreliable, weak, and unlikely to have a strong impact on downstream visual processing. However, our core analyses treat each LGN recording independently and do not consider whether a more reliable signal emerges at the population level. Ideally, this question would be addressed using large numbers of simultaneously recorded LGN neurons. Unfortunately, our dataset consists of many sequentially recorded single units, which limits our ability to study true population dynamics.

Nevertheless, by constructing pseudo-populations from our data, we can ask whether the attentional signal becomes more robust when LGN neurons are considered collectively. Specifically, we asked whether an ideal observer (e.g., downstream neurons in V1) could decode the attentional state of a given trial based on LGN population activity. To address this, we used a linear classifier (support vector machine, svm) to test whether trials could be reliably separated into attend-toward and attend-away categories based on pseudo-population activity patterns. Each pseudo-population was constructed by randomly sampling (with replacement) 20 attend-toward trials and 20 attend-away trials from each neuron, resulting in 198 features (neurons) and 40 samples (trials). The svm was trained on this dataset and tested on a separate, non-overlapping validation set constructed in the same way.

Across 10,000 bootstrap iterations, the classifier achieved a mean classification accuracy of ∼75% (Figure 9A; auc = 0.754 +/- 0.001), which was significantly better different than the label-shuffled control (auc = 0.499 +/-0.001; p = 0.0164). The linear classifier performed worse when only parvocellular data (Figure 9B; auc = 0.634 +/-0.001, shuffle auc = 0.500 +/-0.001, p = 0.1520) or magnocellular data (Figure 9C; auc = 0.659 +/-0.001, shuffle auc = 0.500 +/-0.001, p = 0.1064) were used. This indicates that attentional modulation was not confined to either the parvocellular or magnocellular pathways, consistent with our single-cell analyses described above. Although both subpopulations performed worse than the full population, this was likely due to the reduced number of neurons in these models; a size-matched control population (containing both parvocellular and magnocellular neurons) also performed worse than the full model (data not shown; AUC = 0.642 ± 0.001). Thus, the pseudo-population analysis suggests that some attentional information is indeed present at the population level in the LGN. We next asked whether this weak signal is broadly distributed across the LGN population or carried by a small subset of neurons. To address this, we first performed a “pooled” SVM analysis in which cell identity was erased. Pseudo-populations were generated as before, but the firing rate values for each feature were randomly drawn from the full pool of trials, independent of which neuron they came from. If the attentional signal was distributed across many neurons, this pooling procedure should have preserved classifier performance. However, it did not; the pooled model performed near chance (Figure 9D; auc = 0.507 ± 0.001), no better than the shuffled control (auc = 0.501 ± 0.001; p = 0.4772). This was also the case when only parvocellular data (Figure 9E; auc = 0.506 +/-0.001, shuffle auc = 0.500 +/-0.001, p = 0.4786) or magnocellular data (Figure 9F; auc = 0.503 +/-0.001, shuffle auc = 0.500 +/-0.001, p = 0.4897) were used. This suggests that the attentional signal is concentrated in a minority of LGN neurons.

**Figure 9.**
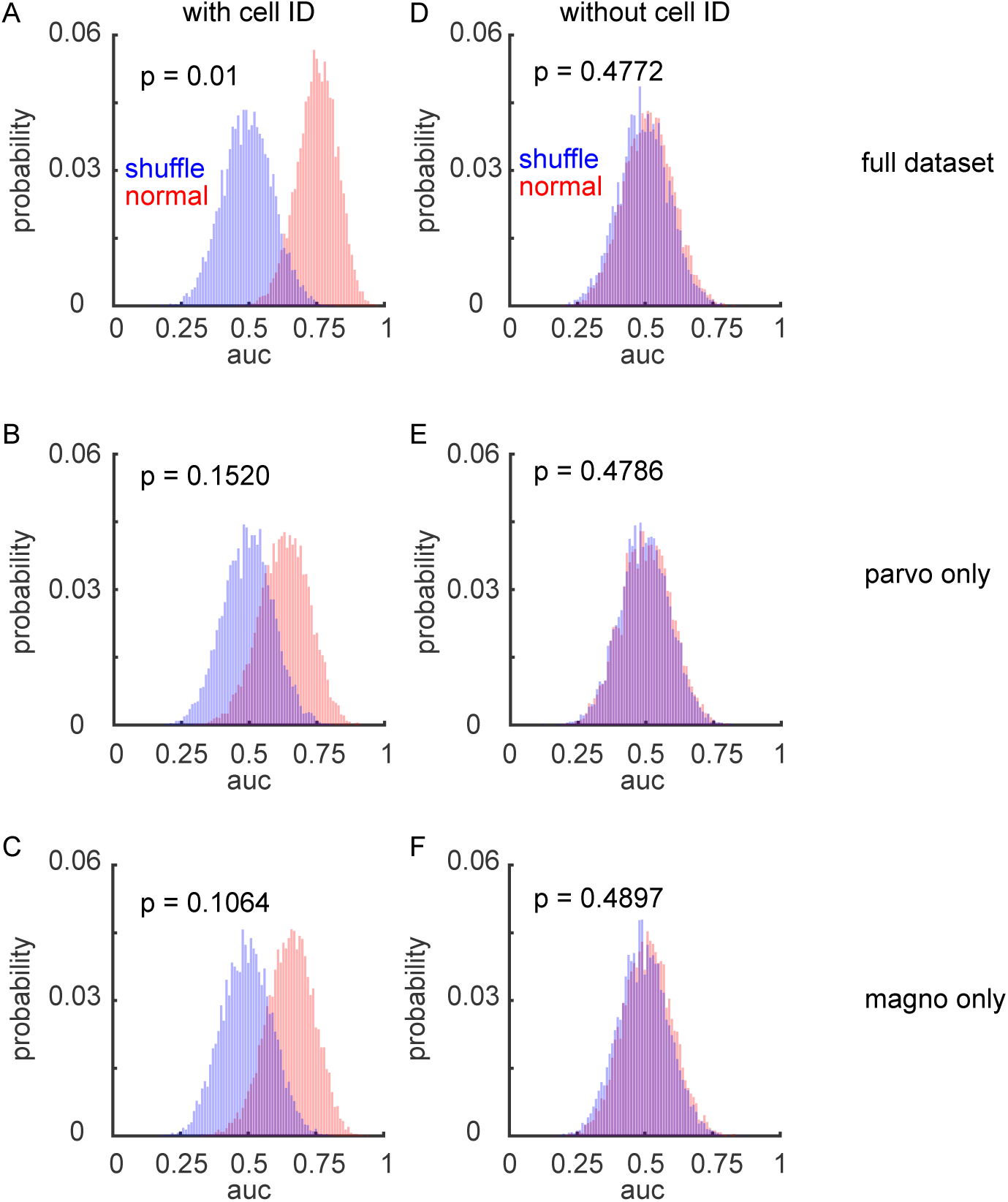
LGN population activity is weakly modulated by spatial attention. Support vector machine (SVM) models were used to assess whether population-level activity in the LGN could predict the subject’s attentional state**. A.** Distributions of classification performance (area under the curve, auc) for svm models trained on pseudo-populations that preserved cell identity (red), compared to shuffled-label controls (blue). **B.** Same analysis as in A, but using “pooled” pseudo-populations in which cell identity was randomized. The sharp drop in performance suggests that the attentional signal is not broadly distributed across the population.

To further test this idea, we applied a “greedy ablation” svm analysis. Here, we began with the full dataset and iteratively removed the most informative neuron, defined as the one with the largest absolute valued beta coefficient, with retraining and re-evaluating the svm after each step. Classifier performance rapidly declined toward chance after the removal of only ∼10% of the neurons (Figure 10A–C), supporting the conclusion that only a small subset of neurons carries the attentional signal. The neurons removed from the model to reach chance discrimination performance were proportionally balanced between the parvocellular and magnocellular pathways compared to random draws from the underlying populations (parvocellular ablations = 14/191, magnocellular ablations = 6/70, p = 0.8155).

**Figure 10.**
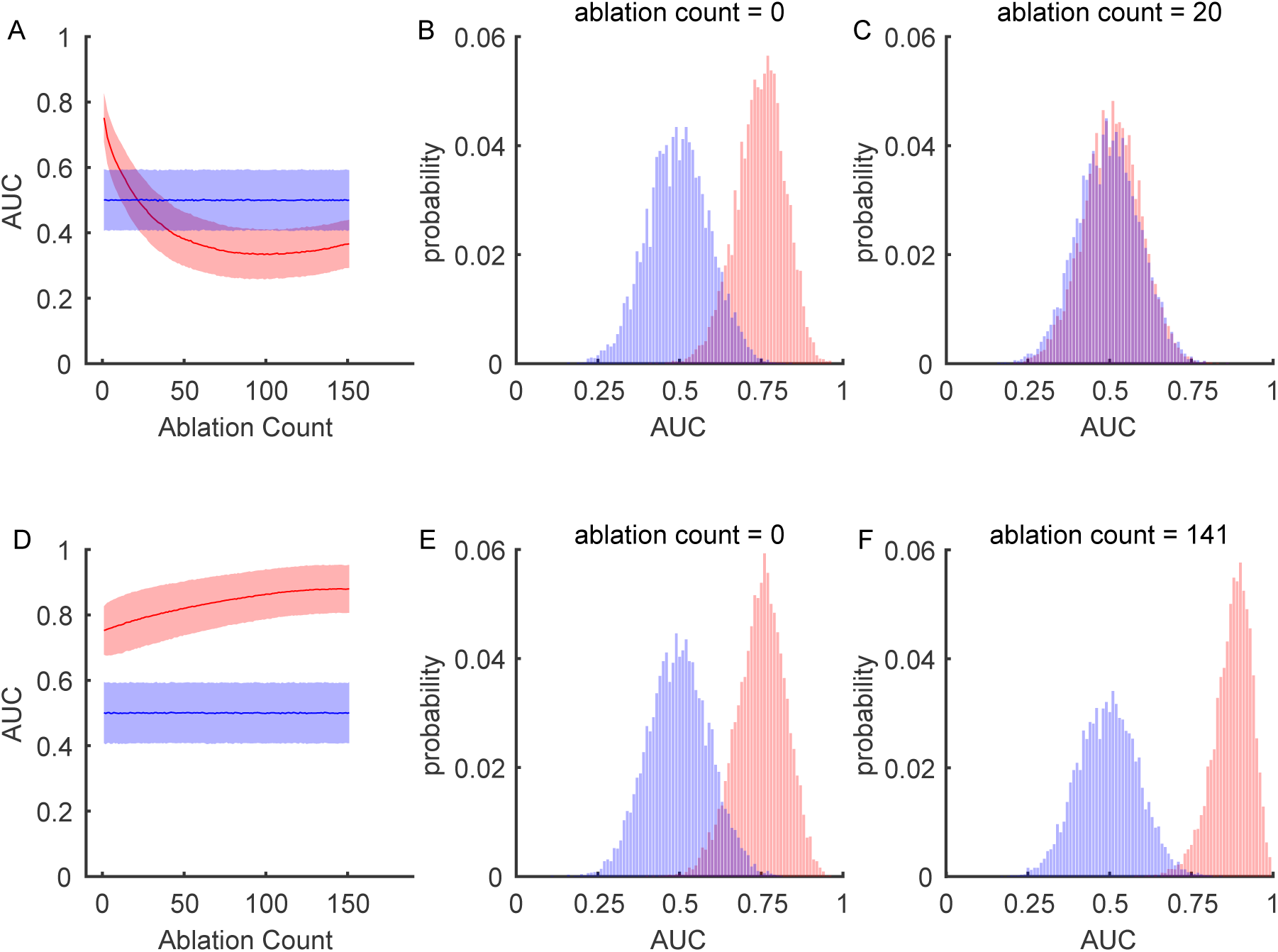
A minority of LGN neurons carry the attentional signal. SVM models were iteratively ablated to assess the relative contribution of individual neurons to the population-level attentional signal. **A.** “Greedy ablation” analysis: the most informative neurons were progressively removed from the model. AUC is plotted as a function of ablation number (0 = full dataset). Shaded areas show ±1 standard deviation across bootstrap iterations. **B.** AUC distribution for the full model (198 neurons). **C.** AUC distribution after 20 greedy ablations, demonstrating a drop to chance performance. **D.** “Reverse greedy ablation” analysis: the least informative neurons were progressively removed. Classifier performance increased as uninformative neurons were removed, peaking with ∼18% of the neurons remaining. **E.** AUC distribution for the full model. **F.** AUC distribution after 141 ablations, corresponding to peak model performance.

Conversely, we also performed a “reverse-greedy ablation” analysis, in which the least informative neurons were progressively removed. In this case, classifier performance improved as uninformative neurons were removed, peaking at auc = 0.859 ± 0.001 when ∼18% of the neurons remained (Figure 10D–F). Together, these three complementary approaches strongly suggest that while an attentional signal is present in the LGN, it is not broadly distributed and is instead carried by a minority (∼10–20%) of the neuronal population.

## Discussion

It is generally agreed that the central nervous system lacks the resources to process all visual information that falls on the retina. Instead, the processing of behaviorally relevant information is prioritized at the expense of other, less- or non-relevant visual information. Spatial attention, the preferential processing of visual stimuli at a specific spatial location, is one mechanism that has a profound influence on what sensory information enters conscious perception. While early studies reported moderate attentional modulation in the lateral geniculate nucleus (LGN)— including increased BOLD signal in humans and enhanced parvocellular/magnocellular firing rates in primates (O’Connor et al., 2002; McAlonan et al., 2008) —subsequent findings have indicated otherwise (Jiang et al., 2015; Shah et al., 2022). Notably, Shah et al. (2022) did not find effects of attention in the LGN, despite observing strong attentional modulation in V1. Interestingly, while measures of spatial attention on visual processing within the LGN have been inconsistent using electrophysiology in non-human primates, the effect has been consistently observed in humans using BOLD fMRI analyses (Schneider and Kastner, 2009; Schneider, 2011; Ling et al., 2015; Poltoratski et al., 2019). Possible reasons for these differences are discussed below.

While the results reported here confirm that spatial attention does influence LGN activity (Figures 2 and 6), a deeper analysis of the data indicates that these effects are weak and inconsistent (Figures 3-5 and 9-10). The observed firing rate increases (on average, ∼1 spike/s) corresponded to only a 1–1.8% contrast change, and ROC analysis showed that an ideal observer could barely distinguish attend-toward from attend-away trials (*53% accuracy*). Furthermore, burst firing, a key indicator of thalamic network state, remained unchanged. Finally, using support vector machine analysis of pseudo-populations constructed from our sequentially recorded LGN neurons, we found that attentional modulation was detectable in the population response patterns. However, this signal was limited to a small subset of LGN neurons and did not generalize across the entire sample. Together, these findings suggest that while spatial attention exerts a statistically significant influence on LGN processing, its functional impact is weak.

Given its position between the retina and cortex (Usrey and Alitto, 2015), the LGN might seem like an ideal site for attentional gating. Its small receptive fields could, in theory, support a highly retinotopically precise spotlight of attention, which becomes harder to achieve in higher-order cortex where receptive fields are larger. However, this is not how the visual system operates. Instead, attentional modulation is strongest in higher-order areas (e.g., IT, MT) (Connor et al., 1997; Seidemann and Newsome, 1999; Treue and Maunsell, 1999; Ramezanpour and Fallah, 2022) and weakest at the early stages of processing (Luck et al., 1997; Shah et al., 2022).

One possible reason is that early visual areas serve as a preprocessing hub, establishing a basis set for the encoding of natural images that is robust against changes in basic stimulus properties such as luminance and contrast (Shapley and Victor, 1979; Heeger, 1992; Olshausen and Field, 1996; Alitto and Usrey, 2004; Bonin et al., 2005; Priebe and Ferster, 2012). When visual information reaches the cortex, V1 acts as a distribution hub sending visual information down a variety of parallel pathways, each of which builds upon the relatively simple, spatially compact receptive fields found in the early visual system, ultimately resulting in a unified perception of the visual world (Felleman and Van Essen, 1991). Strong attentional modulation at the LGN stage might introduce undesirable, nonspecific filtering, affecting all downstream visual pathways indiscriminately. A more efficient strategy is to reserve strong attentional modulation for higher-order areas where receptive fields are already task-tuned, for example, MT during motion discrimination or FFA during face recognition. Thus, in the context of spatial attention, the role of the LGN is to maintain a stable, unbiased flow of retinal input to V1, where visual information is distributed across parallel processing streams before attentional filtering occurs at later stages.

A parallel can be drawn to arousal-driven LGN modulation, which globally enhances or suppresses visual signals depending on behavioral state (Molnár et al., 2021; Crombie et al., 2024). Unlike spatial attention, arousal-based modulation affects all LGN neurons indiscriminately, ensuring coordinated gain control rather than interfering with specific spatial and/or feature-processing streams. Similarly, LGN activity is suppressed during modality-specific attention to non-visual tasks, a scenario where broad suppression is beneficial rather than disruptive (Wimmer et al., 2015).

Animal and human experimental models often provide complementary evidence that, when combined, help build a more complete understanding of brain function. In the case of spatial attention and neural processing in the LGN, the two models appear to provide unique insight into how attention affects different components of the neural circuitry. In humans, spatial attention has been reliably shown to modulate LGN activity as measured by BOLD fMRI (Schneider and Kastner, 2009; Schneider, 2011; Ling et al., 2015; Poltoratski et al., 2019). In contrast, similar experiments in nonhuman primates using extracellular recordings to measure spiking activity have yielded far more variable results, ranging from relatively robust modulation to none at all (McAlonan et al., 2008; Jiang et al., 2015; Shah et al., 2022).

The reasons for this discrepancy are unclear, but a key methodological difference lies in the fundamental unit of measurement accessible to each technique. The BOLD signal reflects changes in blood oxygenation and is sensitive to the activity of large neural populations— potentially hundreds of thousands of neurons within a single voxel—averaged over both space and time, which should improve the detection of small changes in firing rate. Importantly, while the BOLD signal is influenced by both spiking and subthreshold synaptic activity, it is only weakly correlated with the spiking output of the underlying network. Thus, it should be viewed as primarily reflecting aggregate input and local processing (Logothetis et al., 2001; Logothetis, 2008).

In contrast, extracellular electrophysiology captures the spiking activity of individual neurons, is largely blind to subthreshold events, and provides a direct readout of the output of the local network to downstream targets. Traditional single-unit recordings, including those used in the present study, typically capture only one or two neurons at a time. Even with modern multi-electrode arrays, the number of neurons recorded simultaneously is orders of magnitude smaller than the population contributing to a single fMRI voxel. Thus, while fMRI provides a population-level view of neural dynamics emphasizing pre-synaptic activity, extracellular recordings offer a highly resolved but sparse sampling of network output. This fundamental difference in spatial scale and sensitivity likely contributes to the apparent inconsistency in results across methodologies.

One potential path toward resolving this discrepancy is to apply analytical methods that characterize population dynamics rather than focusing solely on individual neuron activity. Such an approach, applied to a sufficiently large sample of LGN neurons, may more reliably reveal an underlying influence of spatial attention on LGN processing. Although the current data set is not ideal for this type of analysis (e.g., neurons were not recorded simultaneously), we took steps in this direction by applying support vector machine (svm) classification to our sequentially recorded data. Overall, this analysis partially supports the view that a population code enriches the detectability of spatial attention in the LGN; however, the influence of spatial attention is not evenly distributed across our recording sample. Instead, the svm models suggest that attentional modulation in the LGN is carried by ∼10-20% of the neurons in our sample. This is in line with our finding that 16.6% of LGN neurons had a statistically significant attention index. Of course, this conclusion does not account for possible temporal dynamics in population activity which could depend on how attention influences a variety of factors including network state dynamics (van Kempen et al., 2021; Shi et al., 2022) and noise correlations (Cohen and Maunsell, 2009; Ruff and Cohen, 2014)

If we are to conclude that only a small percentage of LGN neurons are modulated by attention, there are at least three possible scenarios regarding this subpopulation. First, the modulated neurons may correspond to a known LGN cell type, such as inhibitory interneurons or misclassified koniocellular neurons. Second, the modulated neurons may represent a previously unidentified functional subtype. Finally, it is possible that the modulated neurons do not form a distinct category at all. Instead, the attentional signal may be weak, temporally unstable, and only rarely sufficient to drive detectable changes in spiking activity of LGN neurons.

Finally, we would like to address how the results of the SVM analysis should be interpreted in the context of our data. While SVM classification can be informative, it may also miss the functional point of LGN activity during spatial attention. An SVM asks whether an ideal observer could accurately classify attentional state based on the joint activity of an n-dimensional neuronal population (Pooresmaeili et al., 2010). Any neuron whose firing rate is reliably modulated by attention—regardless of whether that modulation is an increase or a decrease—can contribute to building a linear decision boundary in this space. In this sense, both increases and decreases in firing rate are equally useful to the classifier (Hastie et al., 2009).

While SVM classification is a valuable analytical tool, a literal interpretation of the results can be misleading. LGN neurons do not, “encode” attentional state, nor is there reason to believe that downstream areas such as V1 are trying to “decode” attentional state based on LGN spiking activity. The LGN’s functional role is to relay visual information from the retina to the cortex. Spatial attention, likely via non-retinal sources, modulates this relay by slightly prioritizing the transmission of signals that originate from attended regions of visual space. In this view, the attentional state is not encoded in the LGN per se but instead alters the gain or fidelity of visual signals being passed through it.

## Acknowledgements

We thank K.E. Neverkovec, D.J. Sperka and R. Oates for expert technical assistance.

## Notes

Conflict of Interest: Authors report no conflict of interest

Funding sources: This work was supported by NIH grants EY012576, EY036242, and P50MH132642

### Competing Interest Statement

The authors have declared no competing interest.

### Summary of Updates

In the current addition we have added a population analysis through the use of a linear classifier. Although our data represents a large population of sequentially recorded LGN neurons, we generated pseudo-populations and used support vector machines to address the modulation of LGN processing by spatial attention at the population level.

